# From ambiguous object to illusory faces: EEG decoding reveals a dynamic cascade of face pareidolia

**DOI:** 10.1101/2025.09.10.675272

**Authors:** Ziwei Chen, Mengxin Wen, Qinyi Li, Xun Liu, Di Fu

## Abstract

Face pareidolia, perceiving faces in inanimate objects, provides a unique window into how the brain constructs perceptual meaning from ambiguous visual input. In this work, we investigated the neural dynamics of face pareidolia, testing the hypothesis that it emerges from a temporal cascade that integrates global (low spatial frequency, LSF) and local (high spatial frequency, HSF) visual information. With a pareidolia detection task, we combined event-related potentials (ERP), multivariate pattern analysis (MVPA), and representational similarity analysis (RSA) to map the evolving neural code for illusory face perception. Behaviorally, participants experienced pareidolia from both LSF and HSF stimuli, but the percept was strongest for broadband images containing both frequency bands. Electrophysiological results revealed a clear temporal progression. Early processing (<150 ms) was driven by low-level attributes, with distinct P100/N100 components and neural representations for LSF and HSF signals. A critical shift occurred around 145 ms, after which MVPA decoding could reliably distinguish between images perceived as faces and those that were not. Concurrently, RSA showed that the geometry of neural representations shifted to reflect participants’ subjective face-likeness ratings, rather than the initial physical properties of the stimuli. This transition to a higher-order, subjective evaluation was marked by a later N250 component, whose amplitude indexed the cognitive load of integrating the visual cues. Together, these findings demonstrate that face pareidolia is a dynamic neural cascade, from early sensory encoding to higher-order perceptual evaluation, driven by the integration of global configuration and local features of visual stimuli. Our results illuminate how the brain transforms ambiguous input into meaningful social perception, offering insight into the temporal dynamics of visual awareness.

## Introduction

### Face Pareidolia

Face pareidolia, the perception of illusory faces, occurs when individuals detect faces in objects that do not actually contain them (Liu et al., 2014; Palmer & Clifford, 2020; Proverbio & Galli, 2016; Ryan et al., 2016; Wodehouse et al., 2018). These “face-like objects” can be found in both natural and artificial forms (Chalup et al., 2010; Delbaere et al., 2011; DiSalvo & Gemperle, 2003; Martinez-Conde et al., 2015; Romagnano et al., 2024; C. Wang et al., 2022). This perception is typically rapid and automatic, engaging visual mechanisms that are also involved in processing real faces (Caruana & Seymour, 2021; Guillon et al., 2016; Rekow et al., 2022). This predisposition may stem from a visual system highly tuned to detect minimal facial configurations, such as two dots above a line (Ichikawa et al., 2011; Romagnano et al., 2024; Zhou et al., 2021). This sensitivity can lead the visual system to misinterpret ambiguous patterns as faces (Palmer & Clifford, 2020). However, the precise mechanisms governing this process remain debated. Therefore, investigating face pareidolia is crucial for understanding how the brain processes ambiguous visual signals to construct a coherent percept, offering key insights into face perception itself (Atkinson et al., 2018; Palmer & Clifford, 2020; Rahman & van Boxtel, 2022).

The experience of face pareidolia is thought to depend on the interplay between the global configuration of a stimulus and its local features. When people observe face-like objects, they often encounter visual signals that are incomplete and ambiguous (Hull et al., 2023; Taubert et al., 2017; C. Wang et al., 2022). The brain swiftly recognizes these signals by matching the visual input to stored concepts in memory, enabling rapid predictions (Bar, 2007). This phenomenon is likely driven by the brain’s comparison of local facial features in face-like objects with familiar facial templates, which may lead to misinterpretations when there is significant feature overlap (Chen et al., 2023; Hansen et al., 2010; Hull et al., 2023; Rekow et al., 2022). Additionally, studies have demonstrated that the strength of face pareidolia can be predicted by the prominence of local features, such as eye-like elements, in the objects (Ichikawa et al., 2011). Some research indicates that global configuration plays a role in the perception of face-like objects, where simple facial patterns like two dots and a line contribute to the experience of face pareidolia (Chalup et al., 2010; Romagnano et al., 2024; Zhou et al., 2021). Evidence suggests that people focus more on upright face-like images compared to inverted ones (Guillon et al., 2016; Jakobsen et al., 2023), and face pareidolia is more common in upright orientations (Pavlova et al., 2020). This implies that disrupting the global configuration by inverting the image reduces face pareidolia (Bartlett & Searcy, 1993; Schwaninger et al., 2005). However, a critical gap remains in understanding how the brain dynamically integrates these global and local information streams over time to produce the final, coherent perception of an illusory face.

### Spatial Frequency for Pareidolia

To disentangle the contributions of global configuration and local features in face pareidolia, we can use spatial frequency filtering. Spatial frequency describes visual information at different scales: low spatial frequencies (LSF) convey coarse, global shapes, while high spatial frequencies (HSF) carry fine, local details (Devalois & Devalois, 1980; A. Wang et al., 2022). These distinct informational channels are processed by partially segregated neural pathways. The magnocellular pathway rapidly transmits LSF signals, supporting the analysis of global configuration, while the parvocellular pathway conveys HSF signals for detailed feature recognition (Creupelandt et al., 2022; Holmes et al., 2005; Lacroix et al., 2022). This architecture is thought to support a “coarse-to-fine” processing model, where an initial LSF-based global analysis guides the subsequent, more detailed HSF analysis (Bullier, 2001).

This functional distinction is particularly relevant for face perception, where LSF is crucial for processing the global configuration of a face, and HSF is essential for analyzing local features like the eyes and mouth. This processing is supported by a rapid subcortical route that transmits LSF information to quickly detect a face’s overall structure and orient attention (Creupelandt et al., 2022; Hu et al., 2023; Stein et al., 2014; Vuilleumier et al., 2003). In parallel, a slower cortical pathway processes HSF information for the fine-grained analysis of individual facial features (Mitsudo et al., 2011; Sadeh et al., 2011; Taubert et al., 2022). Therefore, we hypothesize that face pareidolia relies on the dynamic integration of these two streams: an initial, LSF-driven analysis of global structure followed by an HSF-driven evaluation of local, face-like features.

### Temporal Dynamics of Pareidolia with EEG

Electroencephalography (EEG) offers the high temporal resolution necessary to track the neural processing of different spatial frequencies during face pareidolia. Specifically, early event-related potentials (ERPs) generated in the visual cortex, such as the P100 and N100, are known to be sensitive to the spatial frequency content of a stimulus (Lin et al., 2022). The P100, a positive-going wave peaking around 100 ms post-stimulus, consistently shows a larger amplitude for LSF than for HSF stimuli (Lacroix et al., 2022; Nakashima et al., 2008; Wang et al., 2023). This effect is not specific to faces and is thought to reflect an early, general-purpose mechanism in the occipital cortex for processing the coarse, global information carried by LSF signals (Beaucousin et al., 2011; Hansen et al., 2012). Following the P100, the occipital N100 component (often termed the N170 in face-selective literature) peaks around 150-170 ms (Santo et al., 2017). In contrast to the P100, this component shows greater sensitivity to HSF signals, reflecting the subsequent analysis of fine-grained local details required for feature identification (Hansen et al., 2012; Rokszin et al., 2016). The subsequent integration of these segregated LSF and HSF streams is reflected in later cognitive ERP components, such as the frontal-central N250. In face pareidolia tasks, the N250 amplitude is typically larger for matched non-face objects compared to images perceived as faces, suggesting it indexes the outcome of a categorical matching process (Proverbio & Galli, 2016). More broadly, the N250 is thought to index cognitive control and conflict monitoring, with its amplitude increasing when the brain must resolve ambiguity or integrate conflicting information (Jiang et al., 2021; Kida et al., 2003; Liu et al., 2012; Tacikowski et al., 2014; Zhang et al., 2022).

### Statement of Problem

Although real faces are often used as benchmarks in visual neuroscience, in the present study we deliberately focused only on pareidolia. Real faces produce robust neural responses that could mask the subtler dynamics of illusory face perception. Focusing exclusively on pareidolia therefore allows us to isolate how the brain constructs meaningful perception from ambiguous input, providing a tractable model for hallucination-like processes. Beyond this methodological rationale, our approach also enables a principled comparison with well-established models of face processing by testing whether the same processing principles apply when no real face is present. In particular, examining the interaction between pareidolia and spatial frequency can reveal whether the “coarse-to-fine” integration observed for real faces is preserved, attenuated, or qualitatively altered in illusory perception.

This study investigates the neural dynamics of face pareidolia by examining EEG responses to face-like objects and matched controls. Using a pareidolia detection task (Uchiyama et al., 2015; Wardle et al., 2020), we manipulated the spatial frequency content of the stimuli to dissociate the contributions of global configuration (LSF) and local features (HSF). Our hypotheses are derived from a hierarchical “coarse-to-fine” processing framework, which posits an initial analysis of basic visual information followed by higher-order perceptual integration.

First, we used ERPs to examine early visual processing, focusing on the occipital P100 and N100 components. Based on the established roles of LSF and HSF in vision (Fleuaris et al., 2008; Halit et al., 2006) and the hypotheses about face pareidolia (Chen et al., 2023; Jakobsen et al., 2023; Rekow et al., 2022; Zhou et al., 2021), we predicted a main effect of spatial frequency on two early components, reflecting a fundamental, pre-categorical encoding of the physical stimulus properties. Specifically, we hypothesized that LSF-containing stimuli would elicit larger P100 amplitudes, while HSF-containing stimuli would elicit larger N100 amplitudes, regardless of the object’s perceived face-likeness.

Second, we investigated the subsequent stage of categorical evaluation, indexed by the frontal-central N250. We hypothesized that this component would reflect the cognitive processing involved in matching the stimulus to an internal face template. Accordingly, we predicted a main effect of stimulus type: larger N250 amplitudes for non-face objects compared to face-like objects, reflecting the greater processing conflict when evaluating an object that mismatches a facial schema (Proverbio & Galli, 2016). Crucially, to test for the integration of information, we also predicted an interaction between stimulus type and spatial frequency on the N250. We reasoned that the cognitive effort required to evaluate an ambiguous, single-frequency image for “face-likeness” would be greater than for an informationally complete image, particularly for face-like objects.

While ERPs can reveal processing stages, they offer limited insight into the evolving content of the underlying neural representations. Therefore, we employed multivariate pattern analysis (MVPA) and representational similarity analysis (RSA) to map this representational timeline. We used MVPA to determine when the brain distinguishes face-like from non-face-like objects, and RSA to determine how the nature of these neural representations evolves over time. We hypothesized a specific temporal progression: neural representations would initially correlate with the stimulus’s physical properties (i.e., spatial frequency) and later shift to correlate with the participant’s subjective face-likeness ratings. This progression would provide direct evidence for the transition from sensory encoding to subjective perception, charting the neural basis of an illusory face judgment (Palmer & Clifford, 2020; Rahman & van Boxtel, 2022).

## Materials and methods

### Participants

An a priori power analysis (G*Power, V3.1.9.7) determined that a sample of 23 participants was required to detect a medium effect size (*f* = 0.25) with 95% power (α =.05) in a repeated-measures ANOVA. We therefore recruited a total of 37 healthy, right-handed adults (19 male, 18 female, 0 non-binary, 0 prefer not to say; *M* ± *SD*_age_ = 22.08 ± 2.62 years, range: 18–29). Data from six participants were excluded: one for failing to follow task instructions and five for performing below 60% accuracy on the embedded catch-trial task. This resulted in a final sample of 31 participants for analysis (14 male, 17 female; *M* ± *SD*_age_ = 22.39 ± 2.64 years, range: 19–29).

### Apparatus and materials

The experiment was programmed, and data were recorded using E-prime 2.0. Stimuli consisted of 120 pairs of images, each containing a face-like object and a matched non-face object, adapted from a previous study (Wardle et al., 2022). Additionally, six sets of face-like object images and matched object images were chosen for the practice phase, and ten sets with a gray cross presented were selected for the test phase. Following a pre-experimental rating procedure (see Supplementary Materials), the original broadband spatial frequency (BSF) images were processed with a second-order Butterworth filter. This filtering created three manipulated versions from each BSF image: 1) HSF images via high-pass filtering (> 2.4 cycles per degree, cpd); 2) LSF images via low-pass filtering (< 1.6 cpd); and 3) LSF & HSF images, created by applying a band-reject filter that removed middle spatial frequencies (MSF; 1.6–2.4 cpd). All resulting images were then normalized for mean luminance and root-mean-square (RMS) contrast using the SHINE toolbox in MATLAB (Willenbockel et al., 2010). Stimuli were presented as 625×625 pixel grayscale images (Figure 1A) on a 21.5-inch Dell monitor (1024×768 resolution, 60 Hz refresh rate) viewed from a distance of 45 cm.

**Figure 1.**
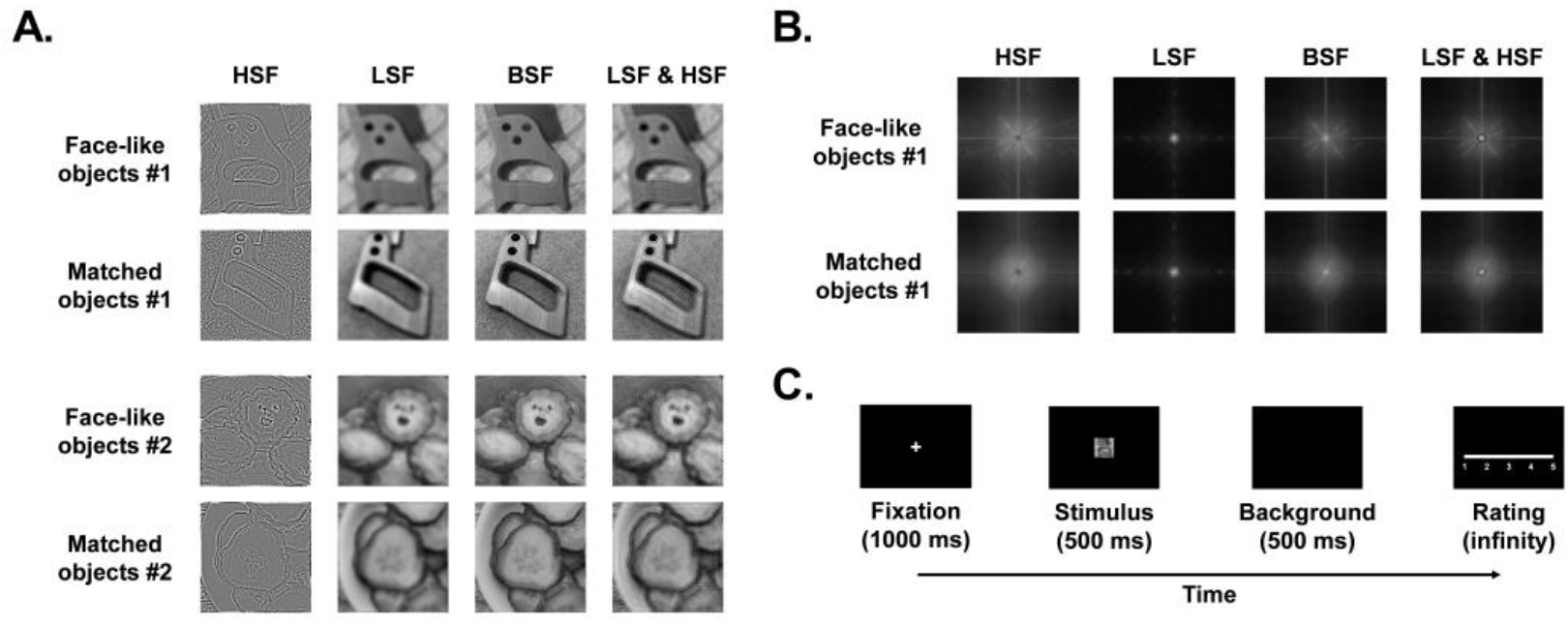
Stimuli and Experimental Design. (A) Example stimuli showing the two types (face-like vs. non-face object) across the four spatial frequency conditions (BSF, LSF, HSF, and LSF & HSF). (B) Example frequency domain plots for BSF stimuli. (C) Schematic of a single experimental trial.

### Procedure

As depicted in Figure 1C, each trial began with a central fixation cross (0.6° × 0.6°) presented for 1000 ms. This was followed by a central stimulus (17.4° × 17.4°) for 500 ms. After stimulus offset, a blank screen was presented for 500 ms, after which participants were prompted to rate the image’s “face-likeness” on a 5-point scale (1 = “very certain no face”, 5 = “very certain a face”). On the 4% of trials that were probe trials, participants were instructed to press the space bar upon seeing the embedded gray cross instead of providing a rating.

The experiment began with a practice phase, followed by the main test phase. The practice phase consisted of 12 trials using stimuli not included in the main experiment, allowing participants to familiarize themselves with both the rating and probe-trial tasks. Participants had to pass a performance criterion on the practice catch trials before proceeding to the main experiment. The test phase consisted of 500 trials presented in random order. This included 480 rating trials (60 trials per condition: 2 stimulus types × 4 spatial frequencies) and 20 probe trials. The entire session lasted approximately 30 minutes.

### Data Analysis

#### Behavioral Data Analysis

Data from six participants were excluded based on pre-defined criteria: five for performing at or below 60% accuracy on the catch-trial task (≥8 errors) and one for non-compliance with rating instructions. The final dataset from 31 participants was analyzed using a 2 (Stimulus Type: face-like objects vs. matched objects) × 4 (Spatial Frequency: HSF vs. LSF vs. BSF vs. LSF & HSF) repeated-measures ANOVA on the mean face-likeness ratings. All post-hoc comparisons were Bonferroni-corrected.

#### EEG Recording and Preprocessing

EEG activity was recorded using a 64-channel cap based on the extended international 10-20 system (Brain Products, Gilching, Germany). The FCz electrode served as the online reference electrode, while the electrode located between FP1 and FP2 served as the ground electrode. Electrodes positioned below the left eye recorded vertical eye movement. The impedance of electrodes was maintained below 10 kΩ, and the sampling rate was set to 500 Hz.

Offline analysis of the EEG data was performed using EEGLAB (Delorme & Makeig, 2004) and custom scripts in MATLAB 2022 (MathWorks, MA, USA). Bilateral mastoid electrodes (TP9, TP10) were utilized as offline reference electrodes, and a low-pass filter at 1 Hz and a high-pass filter at 30 Hz were applied. EEG signals were segmented based on event markers, extracting data from 200 ms before stimulus presentation to 500 ms after presentation, with a baseline of -200 to 0 ms. Trials contaminated by artifacts were rejected, accounting for 0.08% of all trials. Independent component analysis (ICA) was performed, and components related to eye movement, cardiac activity, and muscle artifacts were manually removed. Following preprocessing, data from one participant with poor quality were excluded.

#### ERP Analysis

Based on prior literature and visual inspection of the grand-average waveforms, we defined three components of interest: an early P100 (90–120 ms) and N100 (120–150 ms) over an occipital cluster (PO3, PO4, POz, O1, O2, Oz) and a later N250 (220–260 ms) over a frontal-central cluster (FC1, FC2, F1, Fz, F2). The mean amplitude of each component was then submitted to a 2 (Stimulus Type) × 4 (Spatial Frequency) repeated-measures ANOVA. As with the behavioral analysis, all post-hoc comparisons were Bonferroni-corrected.

#### Multivariate Pattern Analysis

To investigate the temporal dynamics of neural representations, we employed a time-resolved multivariate pattern analysis (MVPA) using a backward decoding approach. The decoding was based on the voltage topographies across all EEG electrodes. To enhance the signal-to-noise ratio and reduce computational demands, the EEG data were first downsampled to 200 Hz and band-pass filtered between 0.5 and 20 Hz (Meng et al., 2023; Tautvydaite & Burra, 2024; B. Zhang et al., 2023). Consistent with common MVPA practices, all trials, regardless of behavioral response accuracy, were included in the analysis. The decoding analysis was implemented using the MVPA-light toolbox (Treder, 2020). For each time point, a linear Support Vector Machine (SVM) classifier was trained to distinguish between trials of different stimulus types (face-like objects vs. matched objects). We used a ten-fold cross-validation scheme to evaluate classifier performance. Within this scheme, all trials were randomly partitioned into ten equally-sized subsets (folds). The classifier was trained on nine folds and tested on the remaining one, with this train-test cycle repeated ten times until each fold had served as the test set. To ensure the stability and reliability of our results against the randomness of data partitioning, this entire ten-fold cross-validation procedure was repeated ten times, each with a new random split of the data. The final decoding performance at each time point was computed by averaging the results across all folds and all 10 repetitions. Classifier performance was quantified using the Area Under the Curve (AUC), where an AUC of 0.5 indicates chance-level performance (Meng et al., 2023; B. H. Zhang et al., 2023). Furthermore, to assess the stability of neural representations over time, we conducted a temporal generalization analysis (Fahrenfort et al., 2018; King & Dehaene, 2014; Li et al., 2022). In this “train-time t, test-time t’“ approach, a classifier trained at a specific time point t was tested on its ability to predict trial labels at all other time points t’. This process was repeated for every time point, resulting in a two-dimensional temporal generalization matrix. The same ten-fold cross-validation and 10-repetition procedure was applied to ensure robust estimates. Significant decoding performance (AUC > 0.5) off the diagonal of this matrix indicates that a neural pattern active at time t is reinstated or maintained at time t’, signifying a stable neural representation. Finally, to investigate the contribution of different spatial frequencies, the entire time-resolved decoding analysis described above was conducted separately for distinct spatial frequency bands of the stimuli.

#### Representational Similarity Analysis

To explore the nature of the neural representations underlying stimulus processing, we employed a time-resolved RSA. This approach quantifies the correspondence between the representational geometry of neural activity and the geometry predicted by theoretical or stimulus-based models. The analysis pipeline consisted of three main steps: (1) constructing neural Representational Dissimilarity Matrices (RDMs) from EEG data, (2) constructing model RDMs based on stimulus properties and subjective ratings, and (3) correlating the neural and model RDMs over time.

For each participant, a time-series of 8×8 neural RDMs was constructed using the same MVPA approach. The neural dissimilarity between any pair of experimental conditions (e.g., condition i and condition j) at a specific time point was defined as the performance of a classifier trained to distinguish between them. The accuracy (ACC) was used as the performance metric, with higher ACC values indicating greater neural dissimilarity between the two conditions. This pairwise decoding procedure was repeated for all 28 unique pairs of the 8 conditions at every time point. The resulting ACC value for each pair was entered into the corresponding off-diagonal cell of an 8×8 symmetric RDM. This yielded a final time-series of neural RDMs for each participant, where each RDM captured the neural representational geometry at a specific moment in time.

Three distinct model RDMs were created to test specific hypotheses about the information encoded in the neural signals. A critical feature of our design was that the specific set of 60 images presented for each condition was randomly drawn for each participant. Consequently, to ensure a valid comparison, all model RDMs were constructed on a per-participant basis rather than being averaged across the group.

Low-Level Attribute Model RDM: This model captured dissimilarities based on low-level visual features. For each participant, we extracted mean luminance and contrast from all presented stimuli, forming a 480×2 feature matrix. A single covariance matrix was computed from this full set of features. We then calculated the average feature vector for each of the eight conditions. The dissimilarity between any two conditions was defined as the Mahalanobis distance between their average feature vectors, using the pre-computed global covariance matrix. This method accounts for the overall distribution of stimulus features specific to each participant.

Spatial Frequency Model RDM: This model tested for representational similarity in the frequency domain. We computed the 2D Fourier transform for each image to obtain its amplitude spectrum. The amplitude spectra for all images within a condition were averaged to create a mean condition-specific spectrum. The dissimilarity between any two conditions was calculated as 1 minus the Spearman rank correlation between their vectorized mean amplitude spectra.

Face-like Rating Model RDM: This model captured dissimilarities based on participants’ subjective judgments. Using each participant’s behavioral data, we calculated the mean rating for each of the eight conditions. The dissimilarity between any two conditions was then defined as the Euclidean distance between their corresponding mean rating scores.

#### Statistical Testing

The statistical significance of MVPA and RSA time-series data was assessed using a non-parametric, cluster-based permutation test (1,000 permutations) to correct for multiple comparisons across time. Clusters were formed by grouping contiguous time points that exceeded an initial statistical threshold (*p* <.05, one-tailed). The sum of the statistical values (e.g., t-values) within each cluster served as the cluster-level statistic. The final significance of each observed cluster was determined by comparing its statistic against the null distribution of maximum cluster statistics, with a cluster-corrected threshold of *p* <.01. For MVPA, this null distribution was generated by randomly flipping the signs of trial labels to test against chance decoding (50%). For RSA, it was generated by shuffling the rows and columns of the model RDM to test against zero correlation (*r* = 0).

## Results

### Behavioral results

First, the scatterplots of mean ratings with different stimulus types and spatial frequencies are shown in Figure 2A. Second, ANOVA revealed a significant main effect of stimulus type, *F*(1, 30) = 1130.493, *p* <.001, *η*_*p*_^2^ =.974, and spatial frequency, *F*(2, 90) = 20.959, *p* <.001, *η*_*p*_^2^ =.411. Additionally, there was a significant interaction between stimulus type and spatial frequency, *F*(2, 90) = 22.847, *p* <.001, *η*_*p*_^2^ =.432. The mean ratings for the eight types of images are shown in Figure 2B.

**Figure 2.**
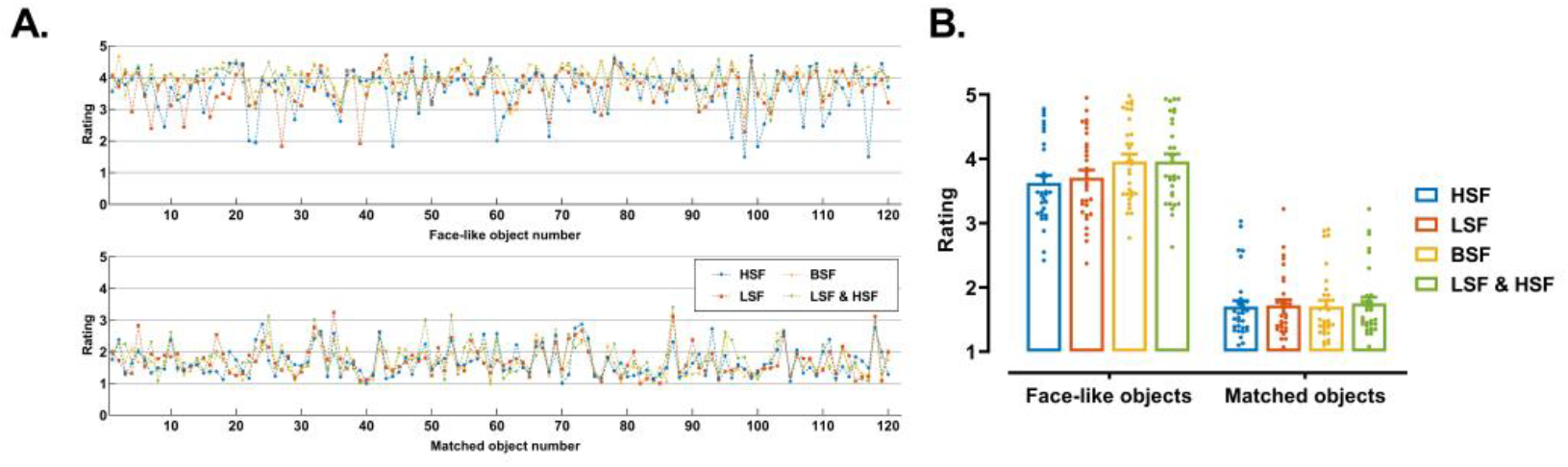
Behavioral Face-Likeness Ratings. (A) Scatterplot of mean ratings for each participant across all conditions. (B) Group-average ratings by stimulus type and spatial frequency. Error bars represent the standard error of the mean (SEM).

Simple effect analyses revealed that in the face-like object group, the ratings for HSF images (*M* = 3.627 ±.65) were significantly lower than those for BSF images (*M* = 3.96 ±.64; *t*[30] = 8.534, *p* <.001, *Cohen’s d* = 1.533) and LSF & HSF images (*M* = 3.96 ±.64; *t*[30] =9.337, *p* <.001, *Cohen’s d* = 1.677). Similarly, the ratings for LSF images (*M* = 3.71 ±.66) were significantly lower than those for BSF images (*t*[30] = 6.820, *p* <.001, *Cohen’s d* = 1.225) and LSF & HSF images (*t*[30] = 6.320, *p* <.001, *Cohen’s d* = 1.135).

### ERP Results

#### Occipital P100

The waveforms on the occipital cluster are shown in Figure 3A. There was a significant main effect of stimulus type, *F*(1, 29) = 10.044, *p* <.01, *η*_*p*_^2^ =.257. Post-hoc tests revealed that the mean amplitude for matched objects (*M* = 1.47 ± 3.38 *μ*V) was greater than that for face-like objects (*M* = 1.08 ± 3.42 *μ*V). Similarly, there was a significant main effect of spatial frequency, *F*(3, 87) = 12.723, *p* <.001, *η*_*p*_^2^ =.305. Post-hoc tests indicated that the mean amplitude for the HSF condition (*M* =.23 ± 4.04 *μ*V) was lower than that for the LSF condition (*M* = 1.52 ± 3.10 *μ*V, *p* =.038), BSF condition (*M* = 1.75 ± 3.15 *μ*V, *p* <.001), and the LSF & HSF condition (*M* = 1.59 ± 3.08 *μ*V, *p* <.001). However, the interaction between material type and spatial frequency was not significant, *F*(3, 87) =.052, *p* =.984, *η*_*p*_^2^ =.002. The mean amplitudes by different conditions are shown in Figure 3B.

**Figure 3.**
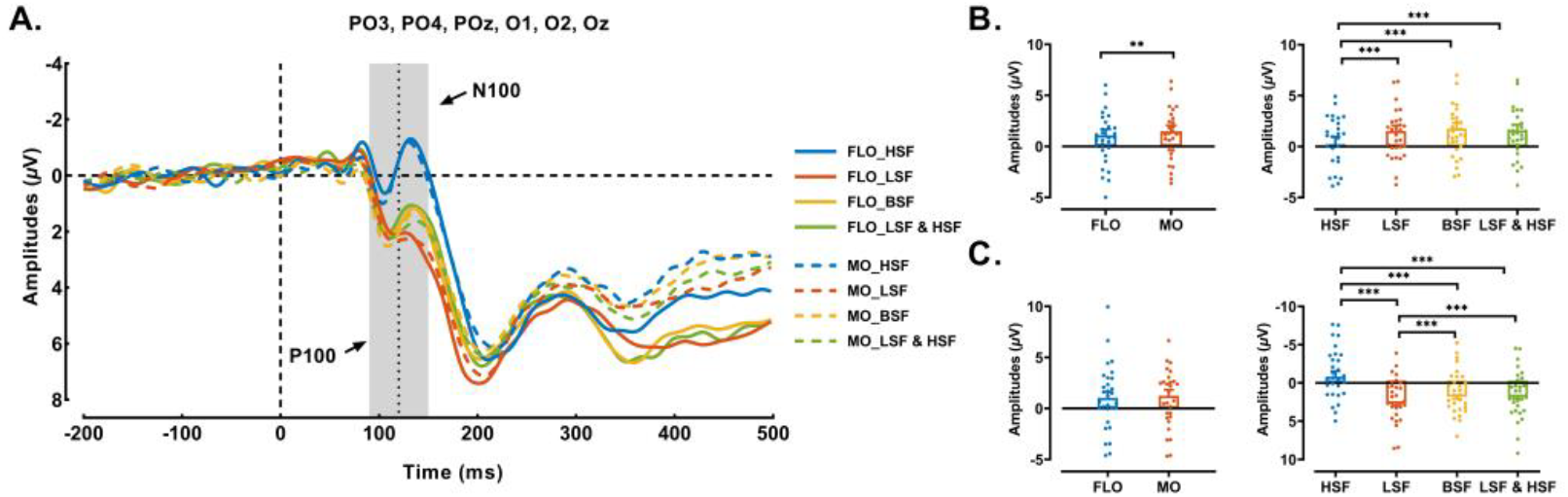
Early Occipital ERPs Show Sensitivity to Spatial Frequency. (A) Grand-average ERP waveforms at the occipital cluster, showing distinct responses to different spatial frequency content. (B) Mean P100 amplitude (90 - 120 ms), showing significant main effects of stimulus type and spatial frequency. (C) Mean N100 amplitude (120 - 150 ms), showing a significant main effect of spatial frequency. Error bars represent SEM. \*\*\**p* <.001, \*\**p* <.01.

#### Occipital N100

The main effect of stimulus type was not significant, *F*(1, 29) = 3.534, *p* =.070, *η*_*p*_^2^ =.109. However, there was a significant main effect of spatial frequency, *F*(3, 87) = 31.633, *p* <.001, *η*_*p*_^2^ =.522. Post-hoc tests revealed that the mean amplitude for the HSF condition (*M* = -.82 ± 3.98 *μ*V) was greater than that for the LSF condition (*M* = 2.38 ± 3.19 *μ*V, *p* <.001), BSF condition (*M* = 1.48 ± 3.27 *μ*V, *p* <.001), and LSF & HSF condition (*M* = 1.52 ± 3.14 *μ*V, *p* <.001). Additionally, the mean amplitude for the LSF condition was significantly lower than that for the BSF condition (*p* =.013) and the LSF & HSF condition (*p* =.040). However, the interaction between stimulus type and spatial frequency was not significant, *F*(3, 87) = 1.096, *p* =.355, *η*_*p*_^2^ =.036. The mean amplitudes by different conditions are shown in Figure 3C.

#### Frontal-central N250

The waveforms on the frontal-central cluster are shown in Figure 4A. There was a significant main effect of stimulus type, *F*(1, 29) = 82.677, *p* <.001, *η*_*p*_^2^ =.740. Post-hoc tests revealed that the mean amplitude for matched objects (*M* = -8.86 ± 4.27 *μ*V) was greater than that for face-like objects (*M* = -6.64 ± 4.62 μV). Moreover, there was a significant main effect of spatial frequency, *F*(3, 87) = 24.910, *p* <.001, *η*_*p*_^2^ =.462. Post-hoc tests showed that the mean amplitude for the HSF condition (*M* = -6.80 ± 4.30 *μ*V) was lower than that for the BSF condition (*M* = -8.47 ± 4.81 *μ*V, *p* <.001) and the LSF & HSF condition (-8.45 ± 4.54 μV, *p* <.001). Additionally, the mean amplitude for the LSF condition (*M* = -7.07 ± 4.52 *μ*V) was lower than that for the BSF condition (*p* <.001) and the LSF & HSF condition (*p* <.001). However, the interaction between stimulus type and spatial frequency was not significant, *F*(3, 87) =.596, *p* =.619, *η*_*p*_^2^ =.020. The mean amplitudes by different conditions are presented in Figure 4B.

**Figure 4.**
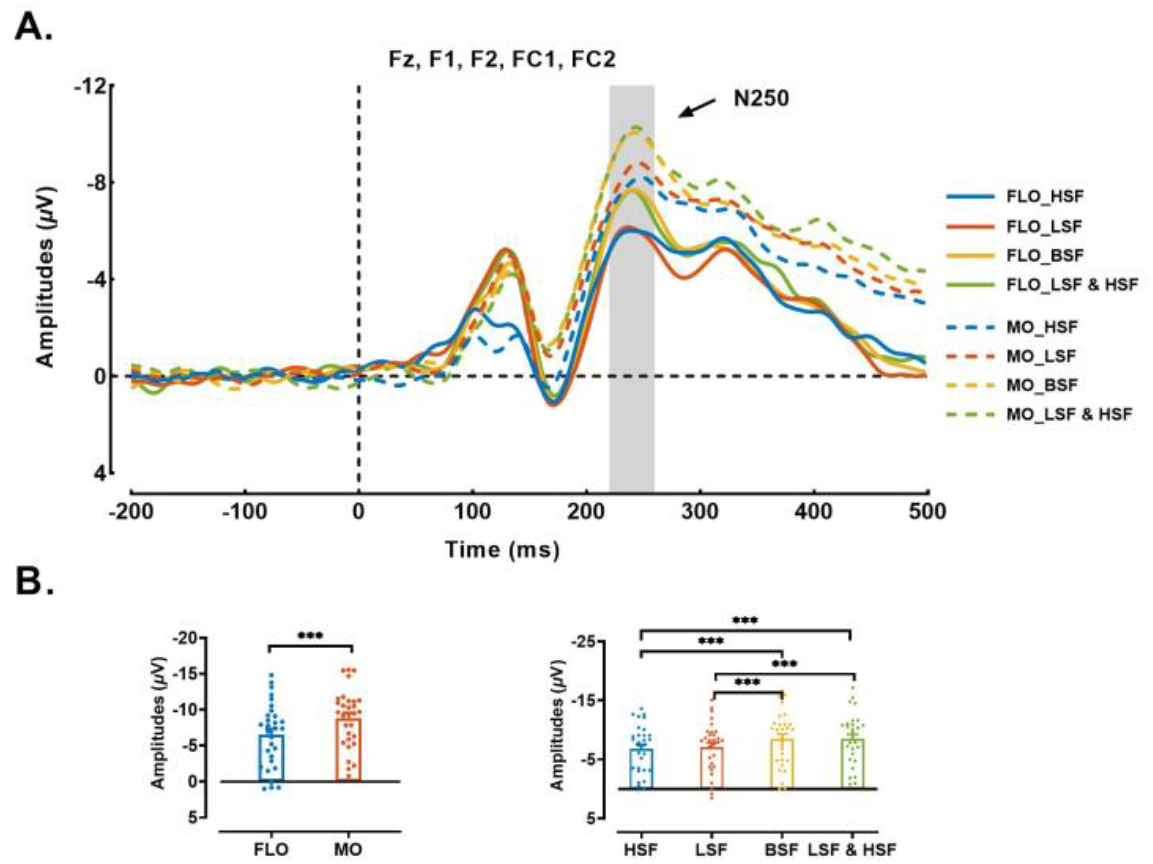
Frontal-Central N250 Results. (A) Grand-average ERP waveforms at the frontal-central cluster. (B) Mean N250 amplitude, showing significant main effects of stimulus type and spatial frequency. Error bars represent SEM. \*\*\**p* <.001.

### MVPA Results

To investigate the temporal dynamics of differentiating face-like objects from matched objects, we conducted MVPA. The results reveal how processing speed and representational stability are modulated by spatial frequency.

#### Temporal Decoding

First, we examined the overall processing timeline by collapsing across all spatial frequency conditions. As shown in Figure 5A, the classifier began to significantly distinguish between face-like objects and matched objects at 145 ms post-stimulus, with this significant decoding persisting until the end of the analysis window at 500 ms (*p* <.01, cluster-corrected).

**Figure 5.**
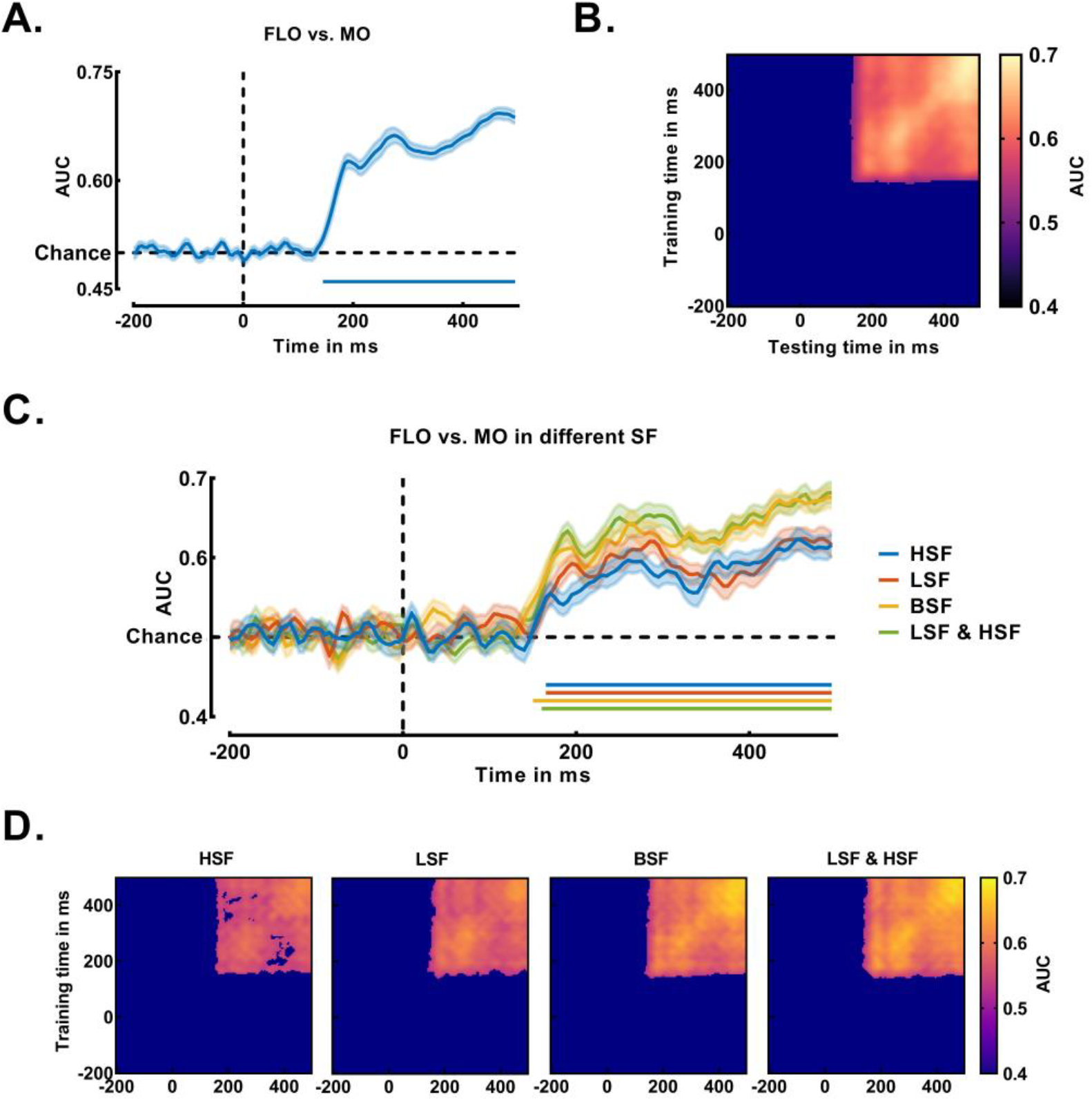
MVPA of EEG data for distinguishing face-like objects from matched objects. (A) Time-resolved decoding accuracy (AUC) for classifying stimulus type, collapsed across all spatial frequency conditions. The shaded area represents the standard error of the mean (SEM), and the solid line along the x-axis indicates time points with significant decoding (*p* <.01, cluster-corrected). (B) Temporal generalization matrix for the classifier trained and tested on data collapsed across all conditions. Saturated colors denote significant generalization performance (*p* <.01). (C) Time-resolved decoding accuracy for classifying stimulus type, shown separately for each spatial frequency condition (HSF, LSF, BSF, LSF & HSF). Colored disks along the x-axis mark significant time points for each condition. (D) Temporal generalization matrices were computed separately for each of the four spatial frequency conditions, illustrating the stability of neural representations within each condition.

To investigate the specific contribution of different spatial frequencies, we then performed the decoding analysis separately for each condition (Figure 5C). This revealed that the neural differentiation occurred earlier for informationally richer stimuli. Significant decoding emerged first for the BSF condition at 150 ms, followed by the LSF & HSF conditions at 160 ms. In contrast, the onset of significant decoding was delayed for single-frequency stimuli, beginning at 165 ms for both LSF and HSF conditions. This pattern suggests that the presence of combined frequency bands facilitates faster categorical processing of the stimuli.

#### Temporal Generalization

To assess the stability and persistence of the underlying neural representations, we conducted a temporal generalization analysis. The overall generalization matrix (Figure 5B), combining all conditions, displayed a large and solid square-like cluster of significance extending from 145 ms to 500 ms. This indicates that the neural patterns distinguishing face-like objects from matched objects were highly stable and sustained throughout the middle and later processing stages.

When examining the generalization matrices for each spatial frequency condition separately, we observed subtle but important differences in representational stability. As shown in Figure 5D, the LSF, BSF, and LSF & HSF conditions all produced robust and largely intact generalization matrices, reflecting highly stable neural representations. In contrast, the generalization matrix for the HSF condition appeared slightly more fragmented, with minor gaps within the significant cluster. This suggests that while HSF information alone is sufficient for the brain to distinguish the stimulus categories, the resulting neural representation may be less stable over time compared to representations built upon LSF or more complete spectral information.

### RSA Results

We compared the EEG RDMs across each time window (Figure 6A) with three model RDMs (Figure 6B). The group-level results revealed distinct time courses for each model (Figure 6C). The Low-Level Attribute model showed a significant correlation with the neural data in two distinct time windows: an early window from 80 to 230 ms and a later one from 345 to 450 ms. The Spatial Frequency model also explained significant variance in the neural data, beginning slightly later and showing a more sustained period of correlation from 105 to 340 ms, followed by another significant cluster from 375 to 495 ms. Finally, the Face-like Rating model demonstrated a robust and prolonged correlation with neural representations, emerging later in the processing stream and remaining significant from 175 to 495 ms. This suggests that subjective judgments are encoded in a stable neural pattern that persists throughout the later stages of stimulus processing.

**Figure 6.**
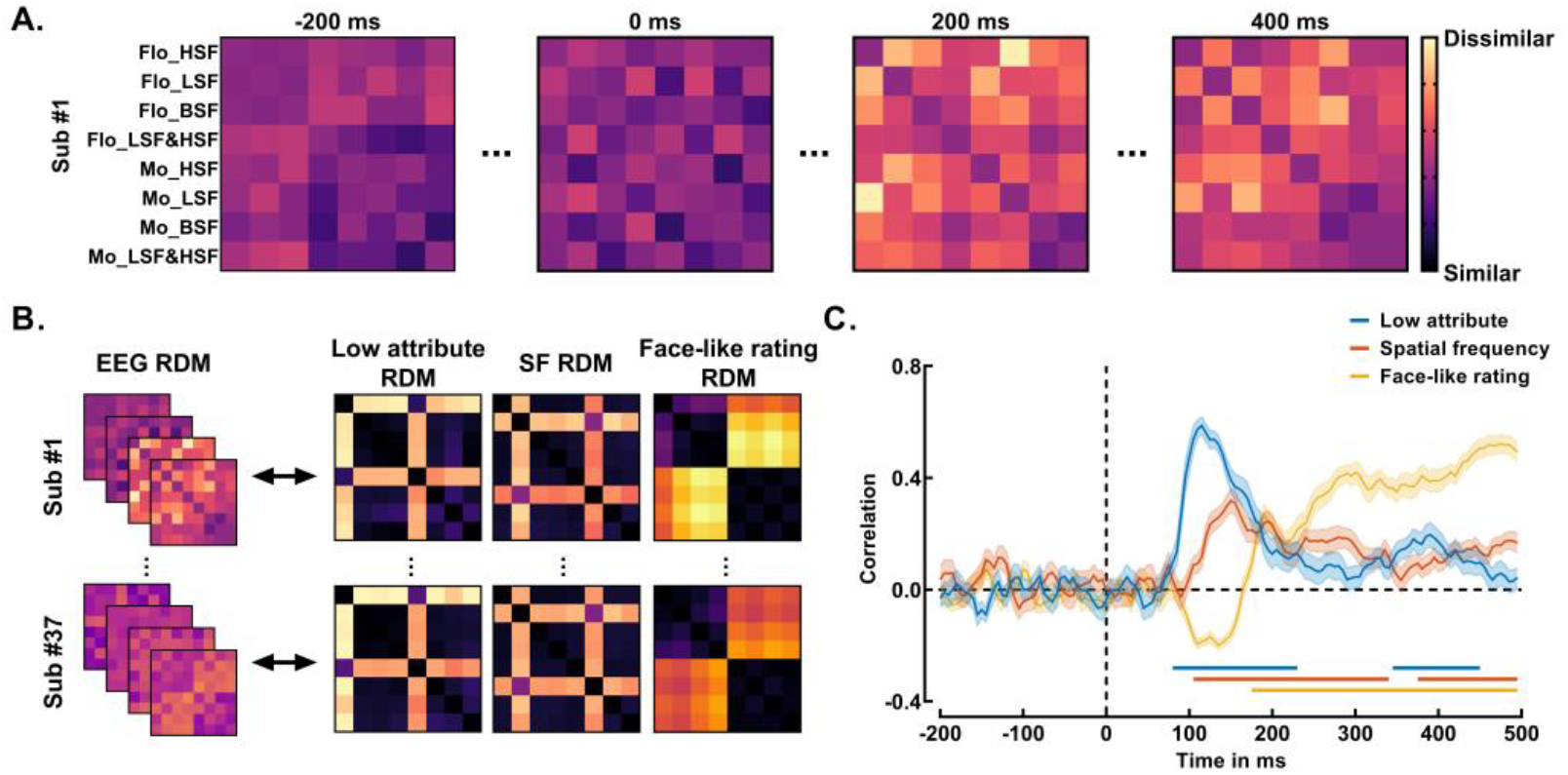
RSA Reveals a Shift from Sensory to Subjective Coding. (A) Example neural RDMs from early (-200 ms) and late (400 ms) processing stages. (B) The three theoretical RDMs used for analysis. (C) Partial Spearman correlations between EEG RDMs and model RDMs for low-level attribute (blue), spatial frequency (red), and face-like rating (yellow). The shaded area represents standard error. Colored disks along the x-axis indicate statistically significant time points.

## Discussion

This study employed spatial frequency filtering to dissociate the roles of global configuration (LSF) and local features (HSF) in face pareidolia. Our behavioral results confirm that face pareidolia is strongest when both LSF and HSF signals are available, supporting the view that the brain integrates global and local information to form the final percept (Chen et al., 2023; Chen et al., 2025; Jakobsen et al., 2023; Rekow et al., 2022; Zhou et al., 2021).

### Interplay of LSF and HSF signals

A central question concerns the relative importance of these global versus local processing streams. In our main experiment, we did not find a significant difference in ratings between face-like images containing only LSF versus only HSF (*t*[30] = 1.743, *p* =.550, *Cohen’s d* = 0.313). However, this null result should be contextualized by our pre-experimental study (*N* = 45; see Supplementary Materials), which revealed a large and significant advantage for LSF images over HSF images (*t*[44] = 6.777, *p* <.001, *Cohen’s d* = 1.010). This robust pre-experimental finding aligns with the classic view that global, LSF-based processing provides the primary scaffold for face perception (Fleuaris et al., 2008; Zhou et al., 2021). The lack of a significant difference in our main experiment may therefore reflect insufficient statistical power for this specific comparison rather than a true absence of an LSF advantage. Taken together, the collective evidence suggests that while both information streams are necessary, global (LSF) processing may play a more foundational role in initially detecting a potential face-like structure. The HSF stream is then critical for analyzing local features, such as eye-like elements, to confirm and strengthen the illusory percept (Ichikawa et al., 2011).

### ERP for LSF and HSF Integration

Our ERP results reveal a clear temporal progression, beginning with early sensory components in the occipital cortex that are sensitive to specific spatial frequencies. Consistent with prior work (Nakashima et al., 2008; Wang et al., 2023), the early P100 component, thought to reflect initial coarse-scale visual analysis, was highly sensitive to LSF content. We found that its amplitude was smallest for HSF-only images and significantly larger for all conditions containing LSF, confirming its role in processing global, LSF-based information. Conversely, the subsequent N100 component is linked to the analysis of fine-grained details and showed greater sensitivity to HSF (Hansen et al., 2012; Rokszin et al., 2016). Accordingly, N100 amplitude was largest for HSF-only images and smallest for LSF-only images, supporting its role in processing local, HSF-driven features. Together, this P100-N100 sequence demonstrates a classic “coarse-to-fine” processing hierarchy in the visual cortex (Hansen et al., 2012), where an initial global analysis (LSF) is followed by a more detailed local analysis (HSF).

Shifting to later, higher-order processing, the frontal-central N250 was sensitive not to the physical properties of the stimulus, but to its categorical interpretation. Specifically, its amplitude was significantly larger for non-face objects than for face-like objects. This aligns with Proverbio and Galli (2016), suggesting the N250 reflects cognitive conflict or a mismatch between a stimulus and a top-down task set, in this case, the search for a face (Zhang et al., 2022). Furthermore, our finding that N250 amplitude was also modulated by the completeness of spatial frequency information underscores the role of the prefrontal cortex as an integration hub. This region must combine context-dependent cues to guide comprehension, a process that can rely on distinct neural routes for different types of information (Amoruso et al., 2023).

### Decoding the Perceptual Timeline

While our ERP results map the initial processing stages, they cannot fully disentangle bottom-up sensory effects from the top-down influence of task expectation. Although the pareidolia detection tasks we used are considered more bottom-up than explicit discrimination tasks (Chen et al., 2023), the instruction to search for a face inevitably creates a top-down expectancy set. To characterize the timeline of this influence, we used time-resolved MVPA. Our results show that a reliable distinction between face-like and non-face objects did not emerge until approximately 145 ms post-stimulus. This timing suggests that a substantial period of bottom-up sensory analysis precedes the formation of a categorical judgment, indicating that top-down expectations may not strongly shape the initial phase of perceptual encoding. Within this timeframe, we found that categorical decoding emerged even later for single-frequency images compared to informationally complete images, suggesting that isolated spatial frequency bands are insufficient for rapid, stable face representation. Furthermore, the finding that the HSF-driven neural representation was less stable over time reinforces the foundational role of the LSF signal in providing the initial structural scaffold necessary for face pareidolia.

Our RSA results provide an even finer-grained view of this timeline, charting the evolution of the brain’s representational geometry from encoding physical features to forming a subjective judgment. This cascade begins with early neural activity (< 150 ms) correlating strongly with the stimulus’s physical properties (our Low-Level and Spatial Frequency models), providing the representational basis for the early P100 and N100 ERP effects. A critical shift then occurs around 175 ms, when the representational structure transitions to correlate strongly and sustainably with the participants’ subjective face-likeness ratings. Crucially, the timing of this shift to a subjective representation (∼175 ms) closely follows the onset of successful categorical decoding in our MVPA (∼145 ms). This convergence of evidence from two distinct multivariate methods strongly suggests that while top-down expectations are critical for pareidolia, their primary influence is exerted in this later time window, after an initial phase of bottom-up feature processing (Bar, 2007; Chen et al., 2023; Hamza et al., 2022; Rekow et al., 2022).

The sustained correlation with subjective ratings, along with the later re-emergence of correlations with physical-level models, strongly suggests the involvement of recurrent top-down feedback (Amoruso et al., 2023; Bullier, 2001; Creupelandt et al., 2022; Stein et al., 2014; Vuilleumier et al., 2003). This suggests a feedforward-feedback loop where higher-order frontal regions send modulatory signals back to the visual cortex to refine the percept, likely by sharpening the representation of HSF details for further analysis in ventral temporal areas (Kauffmann et al., 2015; Mitsudo et al., 2011).

### Limitations and Future Directions

Several limitations of the present study offer clear avenues for future research. First, our stimulus set did not include real faces. Including them would enable a direct comparison between veridical and illusory face perception (Proverbio & Galli, 2016; Taubert et al., 2017; Wardle et al., 2020). Future work could therefore use all three stimulus types to test whether pareidolia recruits an attenuated version of the canonical face-processing network and to directly compare how LSF versus HSF reliance differs between seeing a real and an illusory face. Second, our explicit face-likeness rating task inevitably encourages a top-down search strategy. While our time-resolved analyses suggest this top-down influence emerges relatively late, future studies could better isolate the automatic, stimulus-driven components of pareidolia. For example, using a passive viewing paradigm or a task orthogonal to face judgment (e.g., detecting a rare color change) could reveal the extent to which the perceptual cascade we observed occurs automatically. Finally, our analysis assumes a uniform processing strategy across participants, yet individual traits are known to affect perceptual processing. For instance, autistic traits can modulate the weighting of LSF and HSF information (Alink & Charest, 2020). Additionally, future studies should consider stratifying participants based on their individual sensitivity to pareidolia. Incorporating stable personality or cognitive trait measures may help uncover how such differences shape the neural dynamics of illusory face perception

## Conclusions

In sum, our findings reveal that face pareidolia engages neural representations that closely resemble those elicited by real faces, with comparable temporal dynamics and individual variability. These results suggest that illusory percepts are not mere perceptual noise, but may reflect active top-down inferential processes embedded in the brain’s predictive architecture. Crucially, we identify a transition around 145–175 ms in which neural representations shift from reflecting the physical attributes of stimuli to encoding subjective face-likeness. This turning point represents a candidate neural signature of illusory perception, offering insight into when and how ambiguous input becomes consciously experienced as a face. Additionally, this work lays a foundation for future investigations into the neural basis of hallucination-like phenomena, both in healthy perception and in neurodivergent populations. Going forward, we aim to combine neuroimaging (e.g., fMRI), neuromodulation (e.g., TMS), and computational modeling (e.g., AI hallucination-prone systems) to bridge the gap between human and artificial perception and advance a mechanistic understanding of how hallucinations arise in biological and synthetic minds.

## Data Availability Statement

All data, stimuli, and supplementary materials supporting this study are openly available on the Open Science Framework (OSF): https://doi.org/10.17605/OSF.IO/2J3ER

## Acknowledgements

This work was supported by the National Natural Science Foudation of China (NSFC, No. 62061136001), the Germany Research Froudnation (DFG) under project Transregio Crossmodal Learning (TRR 169). We thank Dr Alan C.-N. Wong, School of Psychology, University of Surrey, for thoughtful discussions and suggestions.

