## Supplementary materials for "From ambiguous object to illusory faces: EEG decoding reveals a dynamic cascade of face pareidolia"

### Methods

#### Participants

A priori power analysis using G*Power (Version 3.1.9.7) indicated that a sample size of 28 participants would be required to achieve a power of 95% for detecting a medium effect size in a two-factor repeated measures ANOVA ($F = 0.25$) at a significance criterion of $\alpha = 0.05$. Accordingly, we recruited a total of 55 participants (21 self-identified as male, 34 as female, 0 as non-binary; $M \pm SD_{\text{age}} = 22.73 \pm 3.05$ years; range 19-30).

#### Apparatus and materials

The experiment was programmed and data were recorded using E-prime 2.0. Based on materials used in previous studies (Wardle et al., 2022), a total of 120 sets of face-like object and corresponding matched object images were selected for the test phase, after screening by four experimenters. Additionally, seven sets of face-like object and matched object images were chosen for the practice phase, and night sets for the test phase. Images with broad-band spatial frequency (BSF) signals were subjected to second-order Butterworth filtering. High-pass filtering (> 2.4 cpd) was employed to obtain HSF images, and low-pass filtering (< 1.6 cpd) was used to obtain LSF images. Subsequently, the SHINE toolbox (Willenbockel et al., 2010) in MATLAB (MathWorks) was utilized to normalize the luminance of all images. All materials were grayscale JPEG files with a resolution of 625 × 625 pixels, as shown in Figure S1A. Participants were seated 45 cm away from a 21.5-inch Dell monitor (resolution: 1024 × 768, refresh rate: 60 Hz).

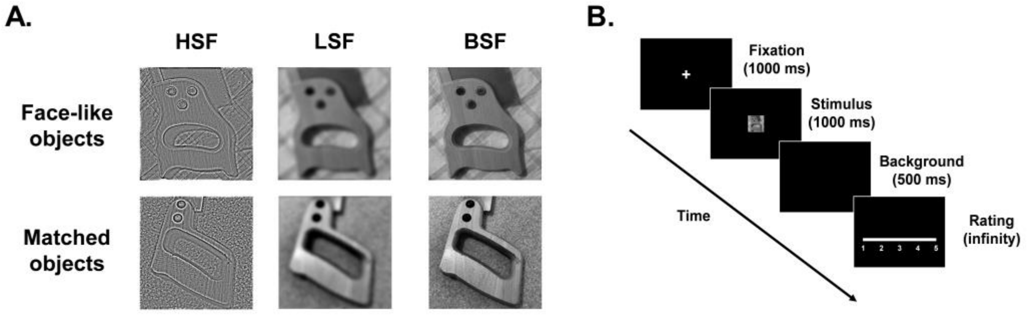


Figure S1. Stimuli and experimental paradigm. (A) Variations of stimuli with different stimuli types (face-like objects vs. matched objects) and spatial frequencies (HSF vs. LSF vs. BSF). (B) The time course of the experimental trial.

### Procedure

The experimental procedure is shown in Figure S1B. In each trial, a fixation cross (0.6° × 0.6°) was presented at the center of the screen for 1000 ms. Subsequently, the material (17.4° × 17.4°) was presented at the center of the screen for 1000 ms. After the material disappeared, a black screen was presented for 500 ms, then a prompt appeared for participants to rate the presence of face-like parts in the image on a scale from 1 to 5 (1 indicating very certain that none existed, and 5 indicating very certain that they existed). Additionally, participants were instructed to press the space bar if they saw a gray cross (0.6° × 0.6°) in the presented image.

The practice phase included 14 trials with 7 sets of images (6 sets for rating trials and 1 set for probed trials) that did not appear in the test phase. Once participants successfully identified 2 probed trials in the practice phase, they proceeded to the test phase. In the test phase, 120 face-like object images and matched object images were used for rating trials, with each image being presented in only one spatial frequency condition (HSF vs. LSF vs. BSF). Each type of material (HSF images vs. LSF images vs. BSF images) was presented in 40 trials. Additionally, 9 sets of images were selected for probed trials, totaling 18 trials. All trial types were presented randomly during the experiment, which lasted approximately 25 minutes.

### Behavioral analysis

Before analyzing the behavioral data, we excluded data from eight participants who made 8 or more incorrect key presses during the 18 probed trials and from two participant who provided arbitrary ratings (reverse scoring) during the test phase, resulting in behavioral data from 45 participants for analysis. First, to investigate whether stimuli type and spatial frequency affect participants' perception of face-like parts in images, we calculated the linear correlation between the mean ratings of the two types of materials at different spatial frequencies. Second, we conducted repeated measures ANOVA with stimuli type (face-like objects vs. matched objects) and spatial frequency (HSF vs. LSF vs. BSF) as within-subject variables and participants' average ratings for each material as the dependent variable. All comparisons were corrected using Bonferroni correction.

### Results

#### Behavioral results

First, the scatterplots of mean ratings with different stimuli types and spatial frequencies are shown in Figure S2A. The mean ratings for face-like object group were consistently higher than for matched object group.

Second, ANOVA revealed significant main effect of stimuli type, $F(1, 44) = 1115.620$, $p < .001$, $\eta_p^2 = .962$, and spatial frequency, $F(2, 88) = 48.129$, $p < .001$, $\eta_p^2 = .522$. Additionally, there was a significant interaction between stimuli type and spatial frequency, $F(2, 88) = 51.593$, $p < .001$, $\eta_p^2 = .540$. The mean ratings for the eight types of images are shown in Figure S2B.

Simple effect analyses revealed that in the face-like objects group, the ratings for HSF images ($M = 3.54 \pm .61$) were significantly lower than those for LSF images ($M = 3.77 \pm .60$; $t[44] = 6.777$, $p < 0.001$, *Cohen's* $d = 1.010$) and BSF images ($M = 4.01 \pm .61$; $t[44] = 12.385$, $p < 0.001$, *Cohen's* $d = 1.846$). Meanwhile, the ratings for LSF images were significantly lower than those for BSF images ($t[44] = 6.028$, $p < 0.001$, *Cohen's* $d = .899$).

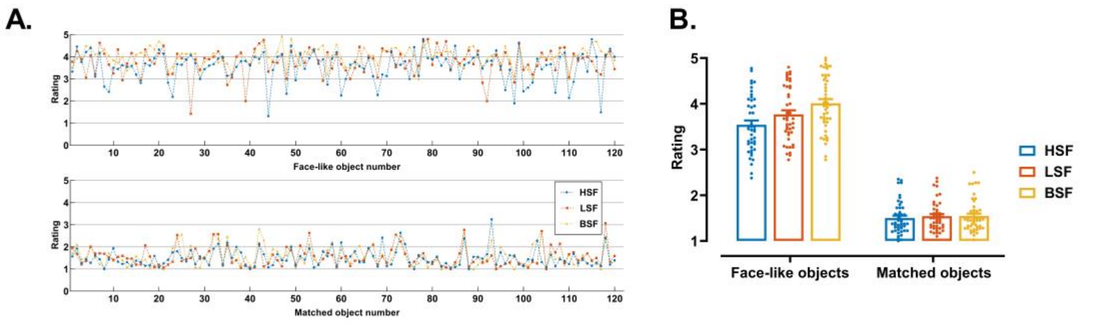


Figure S2. Mean ratings based on the presence of face-like parts. (A) Scatterplot of mean rating with different stimuli types (face-like objects vs. matched objects) and spatial frequencies (HSF vs. LSF vs. BSF). (B) Bar graph of mean rating with different stimuli types and spatial frequencies. Error bars indicate the standard error.

#### Discussion

Among the three levels of spatial frequency, individuals are more likely to perceive face-like parts in face-like objects compared to matched objects. This result indicates that individuals can generate face pareidolia by relying on either LSF or HSF signals of face-like objects. It confirms previous findings that face pareidolia arises from local processing of face-like objects (Chen et al., 2023;

Rekow et al., 2022; Wardle et al., 2020), and shows that the global configuration of face-like objects contributes to this process. Additionally, the intensity of face pareidolia varies among the three types of face-like object images. Individuals rated BSF images highest, followed by LSF images, and HSF images lowest. This result suggests that face pareidolia generated from single LSF or HSF signals of face-like objects is weaker. However, since spatial frequency filtering also removes middle spatial frequency (MSF) signals, differences between BSF images and LSF or HSF images may be due to the absence of MSF signals (Collin et al., 2012; Halit et al., 2006; Wang et al., 2022). Therefore, the differences in ratings among the three types of face-like object images in this pre-experiment may involve the absence of not only HSF or LSF signals but also MSF signals. In summary, the results of the pre-experiment indicate that individuals can generate face pareidolia by extracting either LSF or HSF signals from face-like objects, but they do not clearly determine which type of signal is more relied upon for this process.
